# Ataraxis: Bridging AI Coding Assistants and Scientific Hardware

**DOI:** 10.64898/2026.02.13.705771

**Authors:** Ivan Kondratyev, Weinan Sun

**Affiliations:** Department of Neurobiology and Behavior, Cornell University, Ithaca, NY, USA

**Keywords:** AI-assisted code generation, scientific instruments, Model Context Protocol, laboratory automation, neuroscience, two-photon imaging

## Abstract

AI coding assistants excel at software tasks but lack structured access to laboratory hardware, the physical instruments that define experimental science. We present Ataraxis, an open-source framework that provides hardware control capabilities spanning camera acquisition, microcontroller communication, precision timing, and inter-process coordination, while exposing these capabilities to AI agents through Model Context Protocol (MCP) servers and domain-specific skills. Critically, Ataraxis separates *configuration-time AI assistance* from *runtime data acquisition*, ensuring that experiments run deterministically regardless of AI service availability. We validate this architecture in a two-photon imaging and virtual reality rodent behavior platform, demonstrating up to order-of-magnitude reductions in hardware validation, integration, and personnel onboarding time. By bridging the gap between AI software capabilities and physical instrument control, Ataraxis offers a reusable blueprint for AI-assisted scientific instrumentation across experimental disciplines. All code is available at github.com/Sun-Lab-NBB/ataraxis.

## 1. Introduction

Imagine you are a graduate student preparing for your first brain imaging session. The experimental rig before you includes a specialized microscope capable of recording thousands of neurons simultaneously, a treadmill where a mouse navigates virtual environments rendered across multiple monitors, cameras tracking the animal’s movements, motorized stages positioning sensors with micrometer precision, and micro-controllers orchestrating rewards, timing signals, and sensory stimuli. Before any data collection can begin, every component must be powered, connected, configured, and verified. A single misconfigured parameter or disconnected cable can invalidate hours of experimental work.

You ask your AI coding assistant: “Is the behavior camera connected?” The assistant, capable of writing complex algorithms and understanding vast codebases, can only respond with code to execute. You run the snippet, paste the output, wait for interpretation, receive another snippet, and run again. Ten iterations later, you discover a loose USB cable. A five-second check consumed twenty minutes.

This scenario illustrates a fundamental gap in current AI-assisted development. While tools like GitHub Copilot [1], Claude Code [2], Cursor, and LLM chat interfaces such as ChatGPT [3] have transformed software engineering, scientific research presents unique challenges. Experimental science depends on physical instruments that require configuration, calibration, and troubleshooting. Current AI assistants lack built-in awareness of connected devices, serial port states, or camera capabilities. This forces scientists to context-switch between AI assistance for code and manual debugging for hardware.

The challenge extends beyond simple device queries. Modern scientific instruments generate high data rates [4, 5], demand microsecond-precision timing for synchronization, and require coordination across heterogeneous hardware from multiple vendors. Building and maintaining such systems traditionally requires specialized engineering expertise that academic laboratories struggle to retain. Addressing this demands modular, well-documented libraries that standardize common hardware workflows – camera acquisition, microcontroller communication, precision timing, inter-process data sharing – into reusable building blocks.

In our framework, these libraries form the deterministic runtime layer: the code that directly controls hardware during experiments. But they also serve a second purpose. Because they follow consistent API patterns, provide comprehensive type annotations, and are documented for both human and AI comprehension, they enable AI coding assistants to understand, generate, and compose hardware integration code. To make this practical, we expose hardware discovery and validation through Model Context Protocol (MCP) servers [6] and encode expert workflows as domain-specific skills, creating a structured configuration-time layer through which AI agents help scientists design experiments and write acquisition pipelines. The result is a clean separation (Figure 1): AI assists at configuration time while deterministic libraries handle runtime execution – network outages, API limits, or model errors never affect a running experiment. We call this framework *Ataraxis*, from the Greek *ataraxia* (tranquility), reflecting its goal of undisturbed experimental execution.

**Figure 1.**
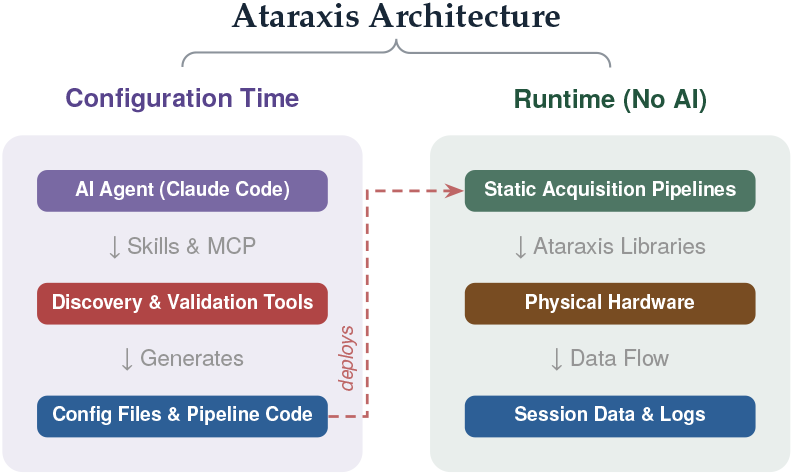
Ataraxis architecture. **Left:** AI agents assist at configuration time via MCP tools and skills. **Right:** Static pipelines run independently during experiments.

We contribute: (1) an ecosystem of 16+ open-source libraries across Python, C++, C#, and MATLAB for camera acquisition, microcontroller communication, precision timing, and inter-process data sharing; (2) MCP servers that expose structured hardware discovery and validation to AI agents; (3) reusable domain-specific skills encoding expert workflows; and (4) data processing and VR environment construction tools. We validate this architecture in a systems neuroscience platform combining two-photon mesoscope imaging with virtual reality rodent behavior experiments (Appendix A, Figure 2) and discuss both opportunities and challenges for AI-assisted scientific instrumentation.

**Figure 2.**
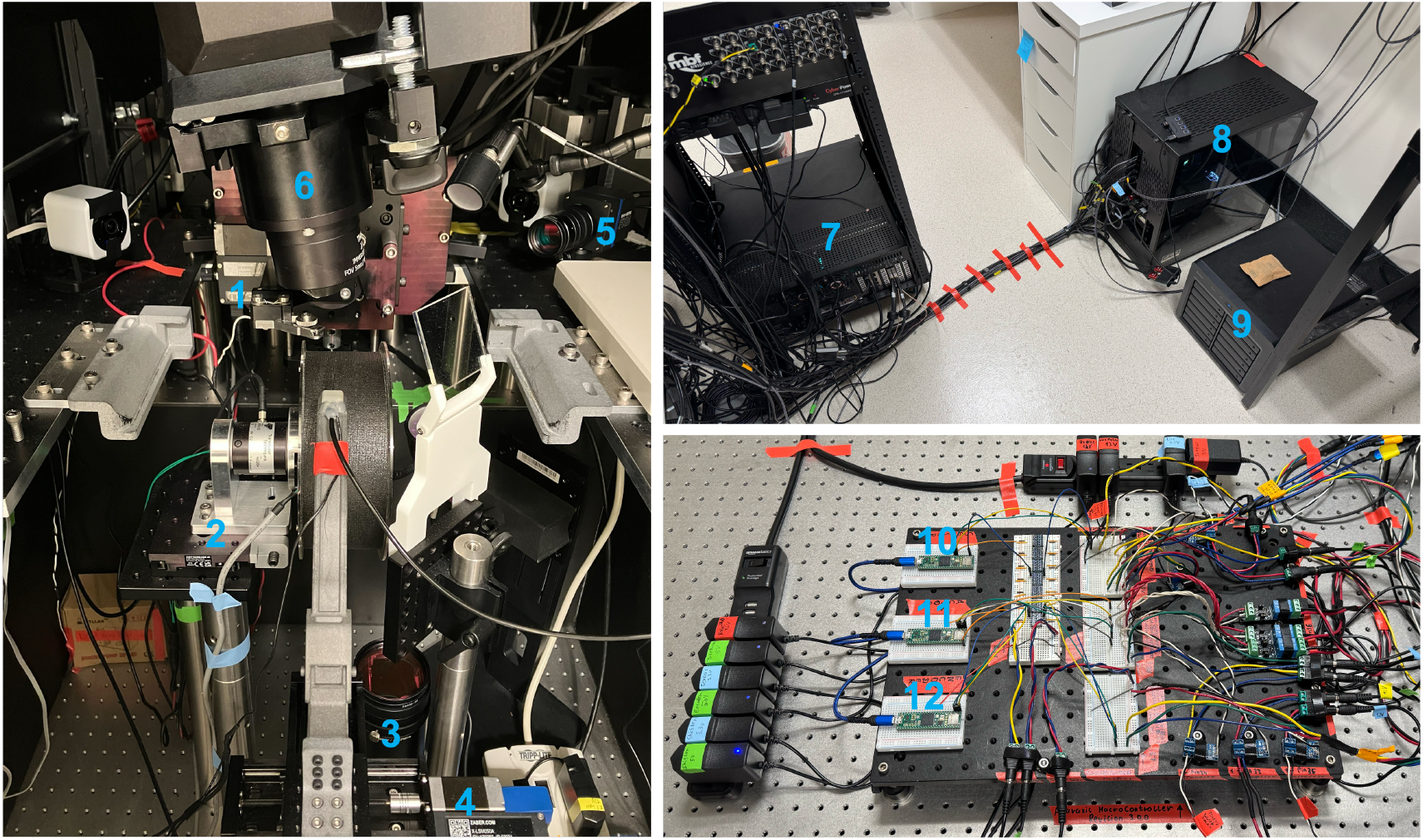
Mesoscope-VR system hardware setup. Labeled components: (1) HeadBar Zaber motor group, (2) Wheel Zaber motor group, (3) FXO546 Camera (face camera), (4) LickPort Zaber motor group, (5) FXO547 Camera (body camera), (6) Two-Photon Random Access Mesoscope, (7) ScanImage PC (Mesoscope controller), (8) Main Acquisition System PC (VRPC), (9) Long-term storage (NAS), (10) Sensor Microcontroller (ID 152), (11) Actor Microcontroller (ID 101), (12) Encoder Microcontroller (ID 203).

Looking ahead, AI agents could move beyond configuration-time assistance to participate in runtime control: writing novel control software, adapting experimental protocols in response to incoming data, and autonomously managing instrument parameters. The comprehensive hardware interfaces that Ataraxis provides today are the foundation for this future. By exposing every relevant control surface through well-documented, typed APIs, the framework ensures that as AI capabilities improve, the transition from assisted to autonomous instrument management can proceed incrementally.

## 2 Related Work

### AI for Scientific Discovery

Recent advances demonstrate AI’s transformative potential across scientific domains. AlphaFold [7] revolutionized protein structure prediction; Sakana’s AI Scientist [8] automates aspects of the research cycle; FutureHouse’s Kosmos [9] pursues autonomous scientific discovery; and Google’s AI Co-Scientist [10] assists with hypothesis generation. However, these advances focus on computational discovery rather than physical instrumentation. The gap between AI-generated insights and their experimental validation remains bridged primarily by human researchers operating complex instruments manually.

### Laboratory Robotics and Automation

Cloud laboratories [11] and biofoundry alliances [12] enable remote execution of standardized protocols on shared robotic platforms. Software frameworks like PyLabRobot [13] provide hardware-agnostic interfaces for liquid-handling robots, and LLM-driven approaches [14, 15] demonstrate AI-guided protocol execution. These systems optimize *protocol execution* on standardized equipment with well-defined action spaces. Ataraxis targets a different challenge: the flexible *development* of novel, evolving instrument systems where hardware configurations change frequently, new components must be integrated, and the action space itself is under active construction.

### Agentic Instrument Control

Recent work explores giving AI agents direct control over scientific instruments. AILA [16] benchmarks multi-agent LLM systems for automating atomic force microscopy, revealing a “sleepwalking” phenomenon where agents operate outside intended boundaries after minor prompt variations – a key safety concern for runtime instrument control. Gently [17] builds a custom orchestrator for light-sheet microscopy with a sample-oriented interface and layered safety architecture, enabling closed acquisition-analysis loops where vision-language models guide experimental decisions. Ataraxis occupies a complementary design point: by restricting AI to configuration time and leveraging an existing coding assistant, it sidesteps runtime safety risks. The MCP servers and skills that Ataraxis contributes are largely orchestratoragnostic and could be integrated into custom frameworks.

## 3 Framework Architecture

Figure 1 illustrates the core design principle: AI agents assist during development and configuration, while experiments run on static, deterministic pipelines. This separation ensures that network issues, API limits, or model updates never affect data collection.

### 3.1 Hardware Interface Libraries

At the foundation are optimized libraries for camera acquisition, microcontroller communication, precision timing, and interprocess data sharing. These libraries serve dual purposes: they provide building blocks for AI-generated code and form the deterministic runtime layer. Table 1 summarizes the core components.

**Table 1.**
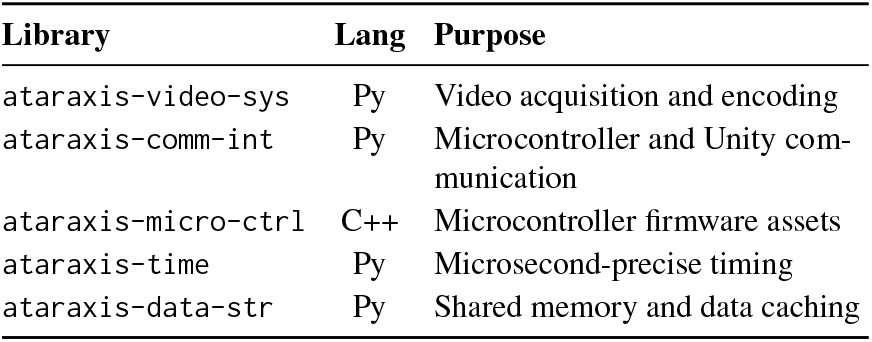
Core Ataraxis libraries. Abbreviations: sys=system, comm-int=communication-interface, micro-ctrl=micro-controller, data-str=data-structures.

The libraries follow consistent API patterns that AI agents can learn and extend. For example, all hardware interfaces expose discovery methods that enumerate available devices, configuration methods that validate parameters before runtime, and acquisition methods optimized for sustained high-throughput operation.

### 3.2 MCP Server Layer

The Model Context Protocol [6] provides structured tool interfaces for AI agents. Unlike shell commands that return unstructured text requiring heuristic parsing, MCP tools return typed data with defined schemas. This enables reliable programmatic processing and, critically, tool discovery: agents can query available capabilities rather than relying on potentially outdated documentation.

Ataraxis implements MCP servers for hardware discovery and configuration validation. When an agent invokes `list_cameras`, it receives structured data including interface type, model, serial number, resolution, and frame rate. When it invokes `check_mqtt_broker`, it receives a clear connectivity status rather than ambiguous terminal output.

Table 2 summarizes the available MCP servers. Full specifications appear in Appendix C.

**Table 2.** Summary of Ataraxis MCP servers. See Appendix C for complete specifications.

| MCP Server | Key Tools | Purpose |
| --- | --- | --- |
| ataraxis-communication-interface | list_microcontrollers,<br>check_mqtt_broker | Discover microcontrollers and test communication channels |
| ataraxis-video-system | list_cameras,<br>check_runtime_requirements | Discover cameras and verify system prerequisites (FFMPEG, GPU encoding, CTI configuration) |
| sl-experiment | get_zaber_devices_tool,<br>get_experiments_tool | Discover additional hardware and list experiment configurations |
| sl-shared-assets | list_available_templates_tool | List VR task templates and shared configurations |

### 3.3 Domain-Specific Skills

Skills package domain expertise as structured instructions that guide AI agents through complex tasks. They address a fundamental limitation: scientific software often spans multiple interdependent repositories with relationships that exceed an agent’s context window.

For example, adding a new camera to the Sun Lab acquisition system requires modifications across four libraries: hardware bindings, configuration schemas, runtime orchestration, and data processing pipelines. The `/camera-interface` skill encodes this cross-repository workflow, specifying which MCP tools to invoke, which files to modify, and which patterns to follow.

Skills can be invoked by name (e.g.,`/system-health-check`) and evolve with the code-base. When hardware changes or best practices improve, updating the skill markdown file immediately updates AI behavior without model retraining. Notably, some skills include self-modification clauses that require agents to update the skill when adding new capabilities, ensuring skills remain perpetually relevant. Table 3 compares Ataraxis capabilities with existing AI coding assistants.

**Table 3.** Comparison of AI coding assistant capabilities. +Ataraxis denotes Claude Code augmented with the Ataraxis framework. 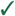 = supported, ° = partial, 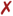 = not supported.

| Capability | Copilot | Cursor | Claude Code | +Ataraxis |
| --- | --- | --- | --- | --- |
| Code completion | ✓ | ✓ | ✓ | ✓ |
| Codebase understanding | ◦ | ✓ | ✓ | ✓ |
| Custom skill-s/rules | ✗ | ◦ | ✓ | ✓ |
| MCP tool integration | ✗ | ✓ | ✓ | ✓ |
| Hardware discovery | ✗ | ✗ | ✗ | ✓ |
| Device status queries | ✗ | ✗ | ✗ | ✓ |
| Pre-session diagnostics | ✗ | ✗ | ✗ | ✓ |

## 4 AI-Assisted Workflows

### 4.1 The Debugging Loop Problem

Consider debugging a camera issue without hardware-aware AI:

**Researcher:** *“The behavior camera isn’t recording*.*”*

**AI:** *“Let’s check if the camera is detected. Run this code*…*”*

*[User runs code, pastes output]*

**Researcher:** *“It says no cameras found*.*”*

**AI:** *“Let’s check the USB connection*…*”*

*[User runs code, pastes output]*

[…*10 more iterations*…]

Each round-trip costs 30–60 seconds and risks transcription errors. Complex systems with multiple hardware components require many such checks, transforming routine validation into an hour-long process.

### 4.2 Hardware-Aware Assistance

With Ataraxis, AI agents assist at two phases: **code generation** using unified libraries, and **pre-experiment validation** using MCP tools. During experiments, static code handles acquisition.

#### Code Generation

**Researcher:** *“I need to add a piezo buzzer for audio cues*.*”*

*[Invokes: /microcontroller-interface skill]*

**AI:** *“Which Teensy should control it? The ACTOR handles outputs*.*”*

**Researcher:** *“ACTOR controller, pin 5*.*”*

*[Generates C++ module and Python bindings]*

**AI:** *“Created buzzer module with pulse, toggle on, and toggle off commands. Want me to upload the firmware and update the dependency version?”*

#### Pre-Experiment Validation

**Researcher:** *“Is the system ready for today’s session?”*

*[Invokes: list_microcontrollers, list_cameras, check_mqtt_broker]*

**AI:** *“All 3 microcontrollers responding. Both cameras detected. MQTT connected. No Zaber motors found; check the power supply*.*”*

## 5 Case Study: Mesoscope-VR System

We deployed Ataraxis in the Sun Lab’s primary experimental system: a two-photon random access mesoscope [4] combined with virtual reality behavioral tasks [18, 19] for studying neural circuits in mice.

### 5.1 System Complexity

The two-photon mesoscope is a specialized fluorescence microscope that images calcium activity across the mouse brain, recording thousands of neurons simultaneously. During experiments, a head-fixed mouse runs on a treadmill while viewing virtual reality scenes rendered on surrounding monitors (via Unity over MQTT), navigating virtual tracks and receiving water rewards at designated locations detected via a lick sensor. The system integrates two high-speed GigE Vision cameras (SVS-Vistek FXO546 for face tracking at 120 fps and FXO547 for body tracking at 60 fps, each configured at 2064 ×1544 pixels), three Teensy 4.1 microcontrollers managing reward delivery, lick detection, treadmill encoding, and TTL synchronization signals, seven Zaber motor axes for precise positioning of the headbar, lickport, and wheel assemblies, the mesoscope synchronized with behavioral events via TTL pulses, and a Unity-based VR environment communicating over MQTT. Appendix A and Figure 2 provide detailed hardware specifications and system photographs.

Even routine maintenance of such systems – replacing a camera, updating firmware, or reconfiguring motor positions – involves coordinating changes across multiple codebases and hardware interfaces. Ataraxis addresses this by encoding engineering knowledge into libraries and skills that persist beyond individual personnel and remain accessible to AI agents.

### 5.2 Deployment Results

Table 4 summarizes estimated improvements observed during Ataraxis deployment. Pre-session validation, which previously required manually checking each hardware component through separate terminal commands, now completes through a single natural language query. Beyond simply diagnosing problems (e.g., “camera is not online”), the AI-assisted approach preemptively offers solutions (e.g., “check if camera is powered”). Adding new hardware interfaces, which typically required days of reading documentation and writing boilerplate code, now takes hours with AI-guided code generation.

**Table 4.** Time comparison for common tasks before and after Ataraxis deployment. Estimates based on lab experience over 6 months.

| Task | Before | With Ataraxis |
| --- | --- | --- |
| Pre-session hardware check | 20+ min | <1 min |
| Debug camera connectivity | 15–30 min | 2–3 min |
| Add new microcontroller module | 2–3 days | 3–4 hours |
| Create experiment config | 30–60 min | 5–10 min |
| Onboard new lab member | 2–3 weeks | 3–5 days |

**Table 5.** Mesoscope-VR system hardware components and interface libraries.

| Component | Hardware | Interface |
| --- | --- | --- |
| Behavior cameras | SVS-Vistek FXO546/FXO547 | ataraxis-video-system (GigE Vision) |
| Microcontrollers | Teensy 4.1 ×3 | ataraxis-communication-interface (USB Serial) |
| Motor positioning | Zaber X-LSM controllers with assorted motors (7 axes) | sl-experiment (zaber-motion Python bindings) |
| VR environment | Unity 6.3 LTS + 3 monitors | sl-unity-tasks (MQTT via ataraxis-communication-interface) |
| Neural imaging | Two-photon mesoscope | ScanImage MATLAB integration |

Ataraxis provides three categories of assistance: **Pre-session validation** verifies hardware connectivity and configuration through MCP tools. **Experiment configuration** guides researchers through template selection and parameter customization. **Hardware interface development** accelerates adding new components by generating code following established patterns.

Video demonstrations and representative interaction transcripts are available in Appendix B.

## 6 Adoption Roadmap

The framework separates reusable infrastructure from lab-specific implementation. Core `ataraxis-*` libraries are available via PyPI; labs create their own ecosystem combining these with custom configurations and skills.

The Sun Lab’s `sl-*` libraries serve as open-source templates. Rather than engineering from scratch, AI agents can reference these implementations, copying and adapting patterns for new hardware configurations. This reduces the initial engineering overhead that typically requires months of specialized development.

Adoption proceeds in phases: (1) install core libraries and configure MCP servers; (2) create lab-specific configuration schemas; (3) encode recurring workflows as skills; (4) iterate as hardware evolves. Appendix E provides detailed guidance.

## 7 Challenges and Future Directions

While Ataraxis demonstrates the potential of AI-assisted scientific instrumentation, our deployment revealed several challenges that warrant attention from the broader AI development community.

### Context Window Limitations

Scientific codebases often span multiple interdependent repositories with complex dependency structures. Current AI agents struggle to maintain coherent understanding across these boundaries, sometimes suggesting solutions that work within one library but break integration with others. Emerging approaches such as graph-based code navigation [20], hierarchical context merging [21], and retrieval-augmented generation may help address this limitation.

### Style and Convention Adherence

Despite detailed skill files and explicit instructions, AI agents inconsistently follow nuanced coding conventions. This requires ongoing human oversight to maintain code quality. Improved instruction following and better mechanisms for encoding stylistic constraints would benefit scientific software development.

### Multi-Project Workflows

The current MCP architecture favors single-project contexts. Complex refactoring that spans multiple libraries requires manual coordination to ensure all relevant skills and tools are available. Standardized mechanisms for composing skills across project hierarchies would simplify multi-repository workflows.

### Legacy Code Integration

Many scientific fields depend on legacy software with unclear dependency structures. AI agents are well-positioned to help modernize these codebases, but current context limitations make this challenging. Better strategies for representing and navigating complex legacy architectures would accelerate the transition to maintainable modern implementations.

These challenges are not fundamental barriers but represent opportunities for improvement in agentic AI frameworks. We discuss them in detail in Appendix F.

## 8. Discussion

Ataraxis targets a general pattern: AI assistance for configuration and validation, deterministic execution for data collection. This pattern extends beyond neuroscience. Genomics laboratories managing sequencers, chemistry labs coordinating spectrometers, and physics experiments with detector arrays face similar challenges. Several design patterns serve as reusable templates for other domains: structured hardware discovery via MCP, domain-specific skills encoding expert workflows, consistent API design across hardware families, comprehensive documentation optimized for AI comprehension, and modular code patterns that enable AI agents to transfer knowledge between libraries.

Currently, Ataraxis handles hardware discovery, experiment configuration, and acquisition pipeline generation. Its modular, well-typed architecture is designed so that new hardware primitives – optical alignment, real-time sensor feedback, adaptive stimulus delivery – can be added without architectural changes. As these control surfaces grow, so does the space of experiments that AI agents can help design – and ultimately, AI agents with sufficient hardware access could potentially discover control strategies and performance optimizations that human engineers would not consider. Future work includes standardized instrument schemas for cross-lab interoperability and automated validation of AI-generated hardware interfaces. All code is available at github.com/Sun-Lab-NBB/ataraxis.

## Acknowledgments

We thank the Cornell Mong Fellowship for supporting I.K. We thank Natalie Yeung, Katlynn Ryu, Jacob Groner, and Jasmine Si for their contributions to various Ataraxis subsystems over the past two years, and Chelsea Strawder for testing AI-assisted and algorithmic features. We thank Hari Shroff, Kesavan Subburam, and Magdalena Schneider for their feedback on the manuscript. We thank the Janelia jET team and Spruston Lab members for discussions on various issues including hardware design. We thank Anthropic for developing Claude Code and the Model Context Protocol. This work was supported by Cornell University.

## A Mesoscope-VR Hardware Configuration

This appendix provides detailed specifications of the Mesoscope-VR system hardware used in our case study.

### A.1 Hardware Components

### A.2 System Architecture

## B Video Demonstrations

Video demonstrations of the Ataraxis framework in action are available online. These recordings show actual AI agent interactions with the Mesoscope-VR system through MCP tools and domain-specific skills.

### B.1 Available Demonstrations

**Demo 1: Pre-Session System Validation**. Shows how a simple natural language query replaces multiple manual debugging commands. The researcher asks “Is the system ready for today’s imaging session?” and the AI automatically invokes MCP tools to check all microcontrollers, cameras, and communication channels, providing a comprehensive status report.

**Video:** youtu.be/Ui2AEvFkCoE

**Demo 2: Hardware Troubleshooting & Experiment Setup**. Demonstrates hardware troubleshooting and experiment configuration with AI guidance. The demo covers hardware discovery, diagnosing connectivity issues, experiment creation, and parameter modification.

**Video:** youtu.be/KBgv4zgwwKw

**Demo 3: AI-Guided Hardware Integration**. Shows how AI agents guide the implementation of new hardware modules. The demo illustrates skill invocation, automatic tool selection, and code generation following established patterns for integrating new components into the acquisition system.

**Video:** youtu.be/iemcuTz1_iM

### B.2 Representative Interaction Transcripts

The following transcripts illustrate typical AI agent interactions with the Mesoscope-VR system. Because skills and MCP invocations pass through an AI interpretation layer, actual outputs are inherently non-deterministic; these examples are representative of the system’s behavior rather than exact reproductions.

#### B.2.1 Demo 1: Pre-Session System Validation

**Scenario:** A researcher needs to verify hardware before an imaging session. The following illustrates a typical interaction pattern as it would appear in a terminal or lab interface.

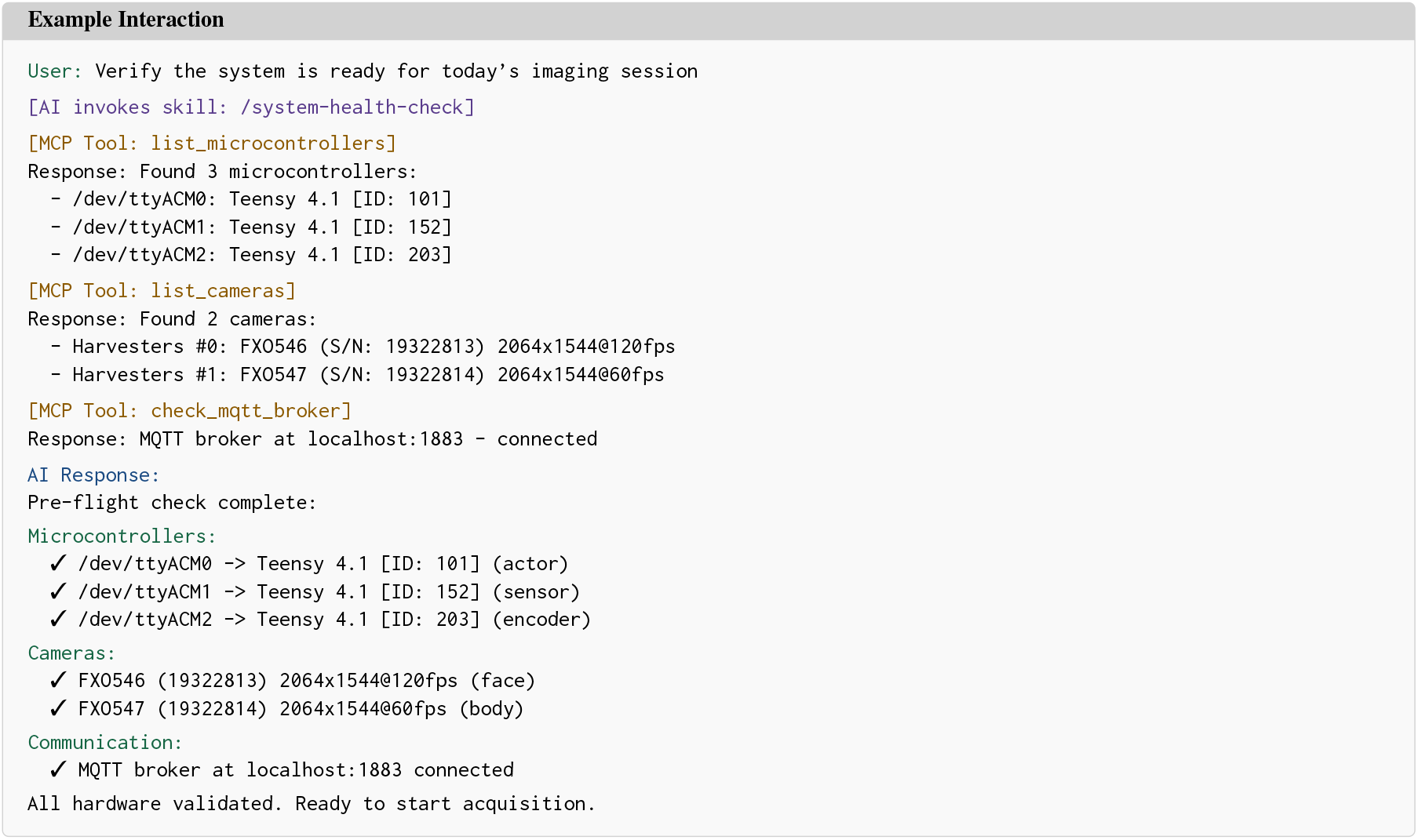

#### B.2.2 Demo 2: Hardware Troubleshooting & Experiment Setup

**Scenario:** Creating a new experiment configuration from a VR task template. This example shows the interactive dialog pattern in a terminal session.

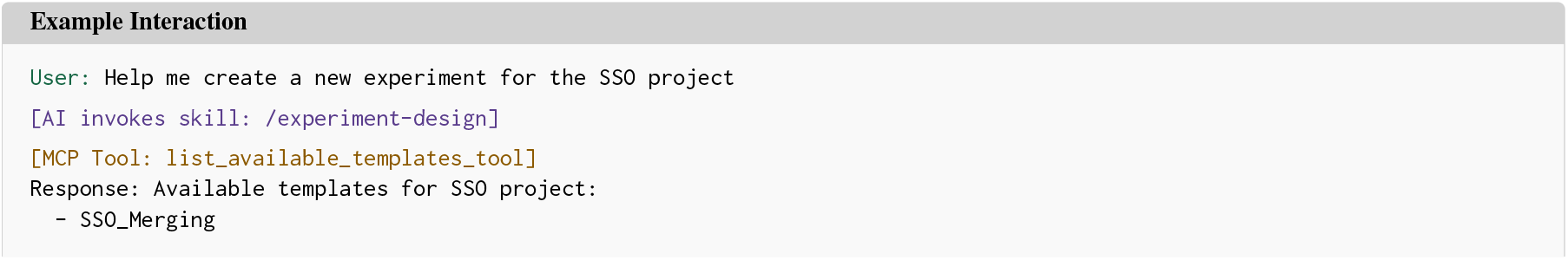

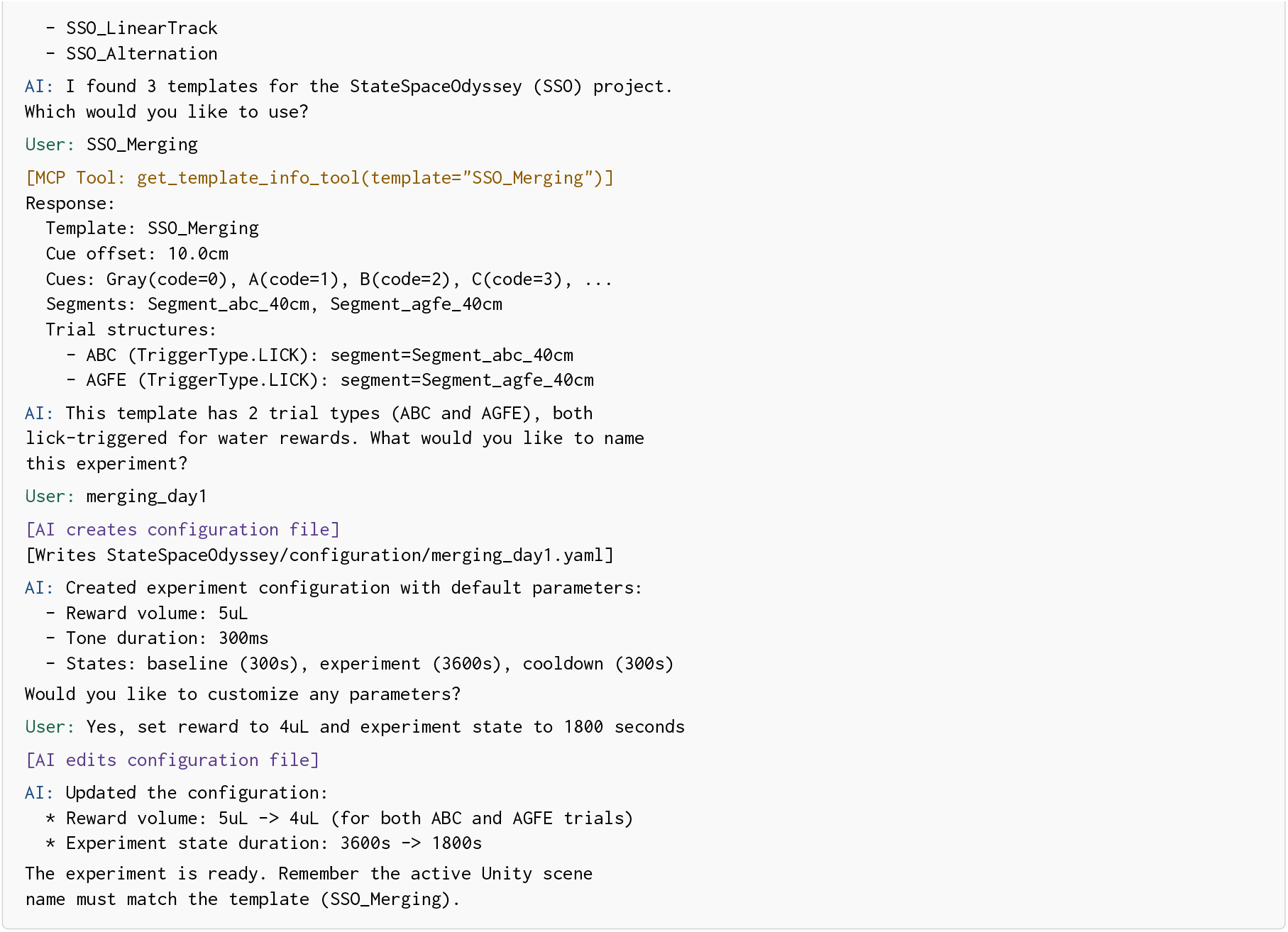

### B.2.3 Demo 3: AI-Guided Hardware Integration

**Scenario:** Adding a new microcontroller module (piezo buzzer) to the acquisition system. This example illustrates the code generation workflow.

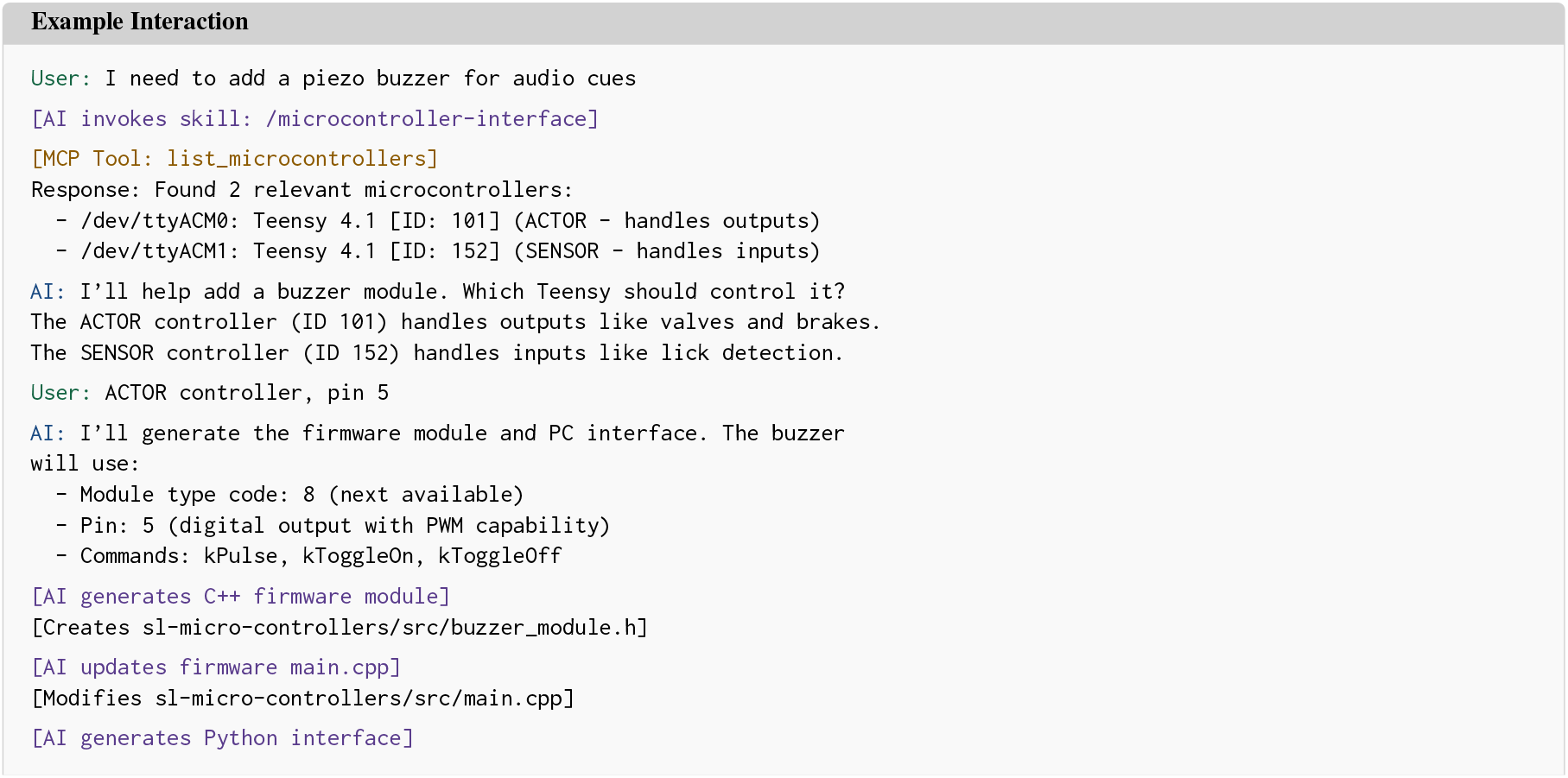

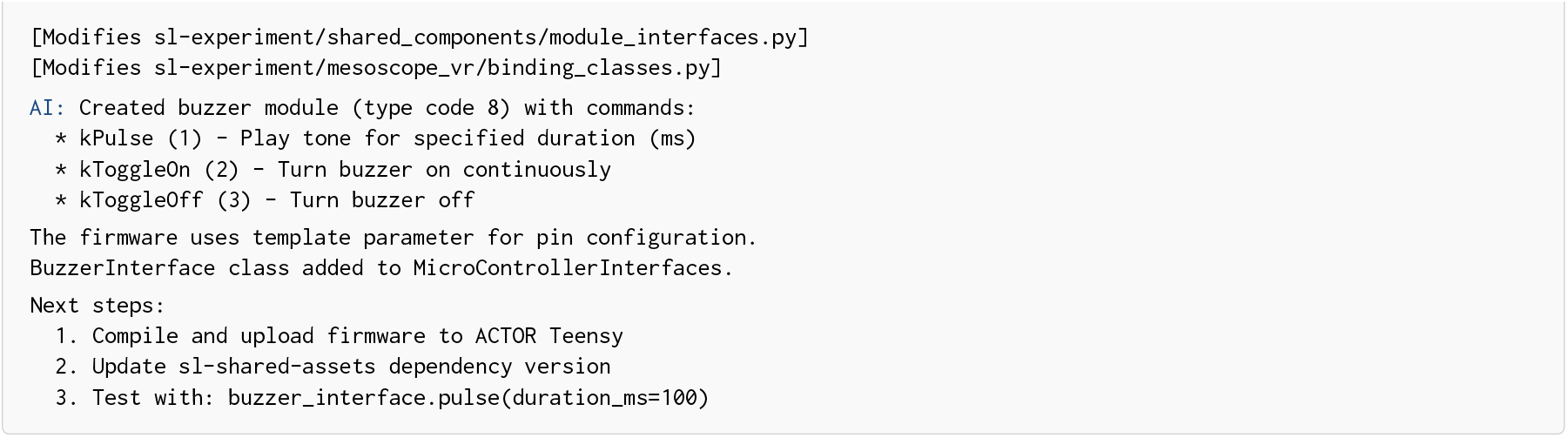

## C MCP Server Specifications

This appendix provides detailed specifications for the Model Context Protocol (MCP) servers implemented in the Ataraxis framework.

Table 6 provides a unified view of all MCP tools available across the four servers. The first two servers (`ataraxis-*`) are reusable across laboratories; the latter two (`sl-*`) are Sun Lab–specific implementations that serve as templates for other groups.

**Table 6.**
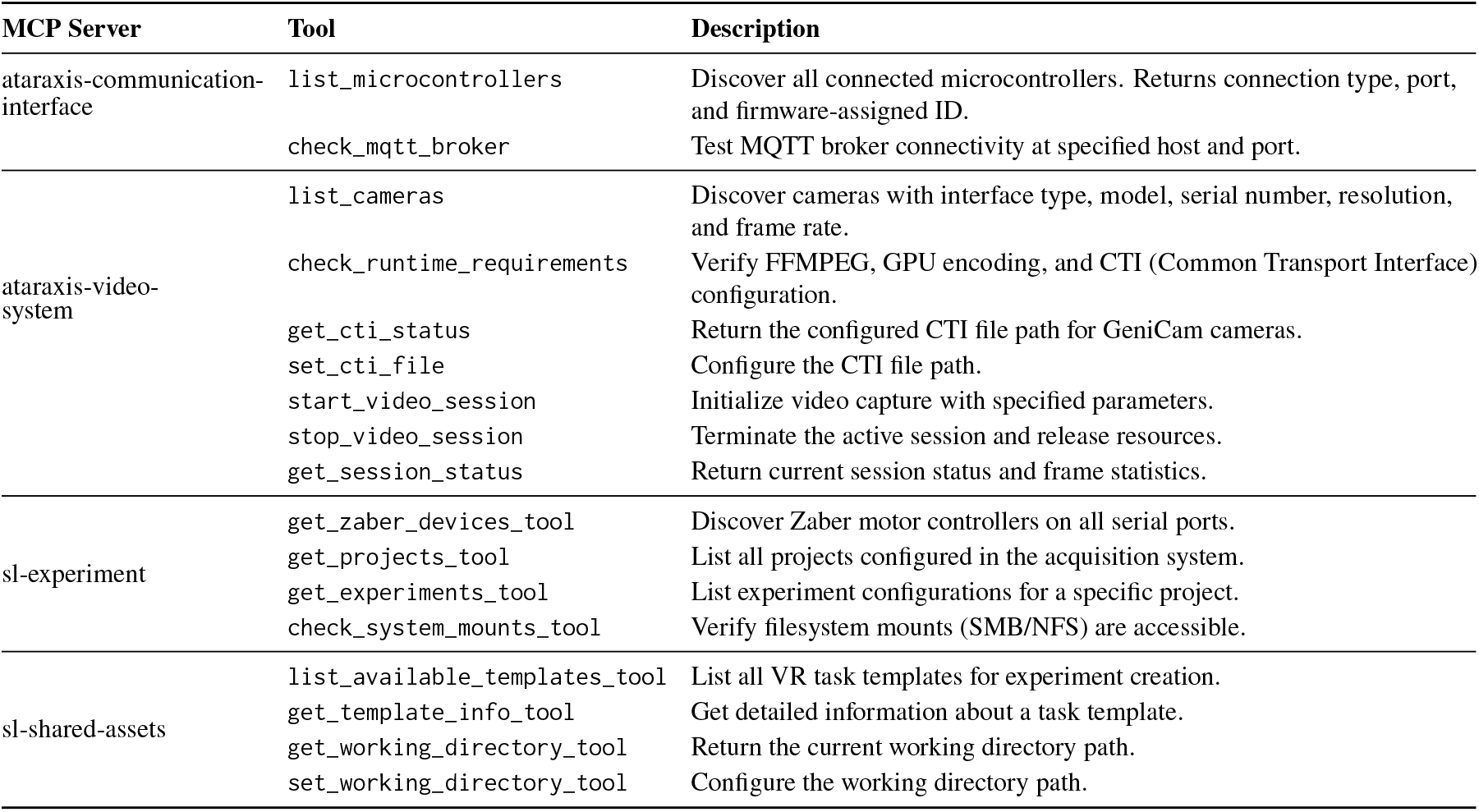
Complete MCP tool specifications across all four Ataraxis servers.

**Table 7.** Domain-specific skills in the Sun Lab ecosystem.

| Skill | Description |
| --- | --- |
| /camera-interface | Guides implementation of camera hardware interfaces using ataraxis-video-system. |
| /microcontroller-interface | Guides implementation of microcontroller modules (C++ firmware) and PC interfaces (Python bindings). |
| /zaber-interface | Guides implementation of Zaber motor interfaces including axis configuration. |
| /experiment-design | Interactive guidance for building experiment configurations using task templates. |
| /system-health-check | Pre-session verification workflow that validates hardware connectivity. |
| /acquisition-system-setup | Guidance for initial system configuration; currently specific to sl-shared-assets. New acquisition systems require nuanced architecture choices best addressed by using sl-experiment as a template with engineer oversight. |

**Combined MCP Configuration** (~/.claude.json):

### MCP Server Configuration

~~~
{
   “mcpServers”: {
     “ataraxis-communication-interface”: {
       “command”: “axci-mcp”,
       “args”: []
    },
    “ataraxis-video-system”: {
       “command”: “python”,
       “args”: [“-m”, “ataraxis_video_system.mcp_server”]
    },
    “sl-experiment”: {
       “command”: “python”,
       “args”: [“-m”, “sl_experiment.mcp_server”]
    },
    “sl-shared-assets”: {
       “command”: “python”,
       “args”: [“-m”, “sl_shared_assets.mcp_server”]
    }
  }
}
~~~

## D Domain-Specific Skills

Skills package domain expertise as structured instructions that guide AI agents through complex tasks.

## E Adoption Roadmap

This appendix provides a detailed guide for laboratories to adopt the Ataraxis framework.

### E.1 Phase 1: Hardware Discovery and Inventory

Begin by using AI agents with Ataraxis MCP tools to discover and document available hardware.

1. **Install A****taraxis** **libraries:**

#### Installation

`$ pip install ataraxis-video-system ataraxis-communication-interface`
2. **Configure MCP servers:** Add Ataraxis MCP servers to your Claude Code configuration file (~/.claude.json).
3. **Run hardware discovery:** Use AI agents to invoke `list_cameras, list_microcontrollers`, and vendor-specific discovery tools to enumerate all connected devices.
4. **Document hardware inventory:** AI generates a hardware manifest documenting device types, ports, serial numbers, and capabilities.

### E.2 Phase 2: Create Lab-Specific Ecosystem

AI agents scaffold the lab-specific library structure using `sl-*` libraries as templates.

1. **Create shared assets library:** Reference `sl-shared-assets` configuration dataclasses as templates for defining your hardware settings.
2. **Create experiment library:** Use `sl-experiment` binding classes as examples for wrapping Ataraxis interfaces with lab-specific defaults and orchestrating data acquisition and preprocessing flows specific to your hardware and infrastructure.
3. **Create microcontroller firmware:** Copy module patterns from `sl-micro-controllers` to implement custom hardware modules.
4. **Add MCP servers:** Follow existing MCP implementations to expose lab-specific discovery and configuration tools.
5. **Write CLAUDE.md:** Document when AI should use MCP tools versus generate code, describing your hardware context and common workflows.

### E.3 Phase 3: Encode Domain Expertise as Skills

Skills capture verified patterns that AI agents use to generate correct code.

1. **Hardware interface skills:** Adapt existing interface skill patterns for your specific hardware.
2. **Experiment design skills:** Use /`experiment-design` as a template for encoding your lab’s experimental paradigms.
3. **System health skills:** Extend /`system-health-check` patterns to validate your specific hardware configuration.

### E.4 Phase 4: Iterate with AI Assistance

Once the foundation exists, AI agents accelerate ongoing development by invoking skills for new hardware, discovering templates for experiments, running health checks before sessions, and updating skills as best practices evolve.

### E.5 Designing for AI Comprehension

The Ataraxis codebase is intentionally structured to maximize AI agent comprehension. All libraries follow consistent naming conventions, provide comprehensive docstrings with typed interfaces, and use uniform patterns across repositories. This consistency enables AI agents to transfer knowledge learned in one library to others: once an agent understands how one microcontroller module is structured, it can generate new modules following the same patterns without additional instruction. Similarly, consistent error handling, configuration schema design, and test organization reduce the cognitive overhead for AI agents working across the ecosystem.

## F Challenges in Agentic AI for Scientific Software

Our deployment of Ataraxis revealed several challenges in using AI coding assistants for scientific instrumentation. We document these observations to inform future development of agentic AI frameworks and to help other laboratories anticipate similar issues.

### F.1 Style Guide and Convention Adherence

The Sun Lab maintains detailed coding style guides to ensure consistency across libraries and to make code more parseable by AI agents. These guides are implemented as skill files following best practices: they use sub-files to stay under recommended length limits, include checklists for AI verification, and are referenced from the CLAUDE.md entry point.

Despite this scaffolding, AI agents inconsistently follow nuanced style prescriptions. For example, our style guide explicitly requires prose in documentation rather than bullet lists, yet agents frequently default to lists. Similarly, commit message formatting often deviates from specified patterns unless the exact phrase “follow the style guide” appears in the prompt.

This behavior persists even in sessions without context compaction, suggesting that the issue is not simply context window overflow. The agents appear to have difficulty maintaining detailed stylistic constraints alongside functional requirements. This creates ongoing overhead: engineers must monitor AI output and correct style violations rather than accepting generated code directly.

**Opportunity:** Improved mechanisms for encoding and enforcing stylistic constraints, perhaps through dedicated style-checking tools or more robust instruction-following capabilities, would reduce this oversight burden.

### F.2 Multi-Project Coordination

The current MCP and skill architecture favors single-project contexts. Project-specific MCP servers, virtual environments, and skill files provide expanded domain expertise for individual libraries. However, many scientific engineering tasks span multiple interdependent projects.

The Sun Lab acquisition system spans four libraries using three languages: a C# Unity library for VR environment control, a C++ library for microcontroller firmware, and two Python libraries for runtime management and data collection. Adding a new experiment configuration requires coordinated modifications across three of these libraries.

When an AI agent is invoked from a parent directory containing multiple projects, it does not automatically discover skills from all sub-projects. Instead, when asked to perform cross-project refactoring, the agent often examines source code directly and proposes solutions that work within one library but break integration with others. For example, rather than using Unity’s GUI-based configuration system (as documented in the Unity project’s skills), an agent might suggest directly modifying serialized object properties, which corrupts the Unity project.

Our current mitigation designates one project as the entry point and manually maps other projects through an elaborate lattice of skills and external MCP references. This approach is fragile and prevents using MCP servers from projects with conflicting dependencies.

**Opportunity:** Standardized mechanisms for composing skills and MCP servers across project hierarchies would enable more natural multi-repository workflows. This might include automatic skill discovery, dependency-aware environment management, or hierarchical context specifications.

### F.3 Legacy Codebase Integration

Scientific computing frequently depends on legacy software that provides validated but unmaintained functionality. These codebases often have unclear or circular dependency structures, outdated language versions, and undocumented APIs. AI agents are theoretically well-positioned to help modernize such code, but current context limitations make this challenging.

We encountered this when integrating the pirt package, which implements diffeomorphic demons registration, into our image processing pipeline. The package had complex circular dependencies that required extensive human guidance to disentangle. The agent could not fit the entire dependency structure into context simultaneously, requiring multiple compacting cycles or deliberate task chunking by supervising engineers.

This challenge likely reflects fundamental limitations in how current AI agents represent and navigate large codebases. Linear context windows struggle with the graph-structured relationships that characterize software dependencies.

**Opportunity:** Graph-based memory representations or alternative architectures for code understanding could significantly accelerate legacy modernization efforts. This would benefit not only neuroscience but many scientific fields that depend on aging but validated software, from climate modeling to genomics analysis pipelines.

### F.4 Summary

These challenges are not fundamental barriers to AI-assisted scientific instrumentation. Rather, they represent areas where current agentic AI frameworks have room for improvement. We share these observations in the hope that they inform the development of more capable tools for scientific software engineering.

The Ataraxis framework demonstrates that meaningful AI assistance is already possible within these constraints. By carefully designing skills, structuring codebases for AI comprehension, and maintaining human oversight for quality assurance, laboratories can substantially accelerate their instrument development workflows.

